# A First-in-Class Dual Degrader of Bcl-2/Bcl-xL Reverses HIV Latency and Eliminates Ex Vivo Reservoirs

**DOI:** 10.1101/2024.10.16.618723

**Authors:** Lin-Chun Chang, Michael T Yin, Gregory M. Laird, Kristen D. Ritter, Jayesh Shah, Asim K. Debnath

**Author notes:** Correspondence: Lin-Chun Chang; Asim K. Debnath Laboratory of Molecular Modeling and Drug Design, Lindsey F. Kimball Research Institute, New York Blood Center, New York, New York.

## Abstract

The persistence of latent HIV-1 proviruses in CD4+ T cells is a major obstacle to curing HIV. The “shock and kill” strategy involves reversing latency with latency-reversing agents (LRAs) and selectively inducing cell death in infected cells. However, current LRAs have shown limited efficacy in eliminating the ex vivo HIV reservoir. We repurposed PZ703b, a pro-apoptotic protein degrader initially developed for anti-leukemia therapy, to target HIV eradication. PZ703b induced degradation of Bcl-2 and Bcl-xL, activating the non-canonical NF-kB pathway and caspases cascade, resulting in latency reversal and selective apoptosis of infected cells. Treatment of ex vivo CD4+ T cells from ART-suppressed HIV-1 patients achieved a ∼50% reduction in the replication-competent reservoir. Our study provides proof-of-concept for using protein degraders to reverse HIV latency and induce cell death, highlighting PZ703b’s potential in HIV cure strategies. This approach may pave the way for novel therapeutic interventions aimed at eliminating the HIV-inducible reservoir.

## Introduction

Combination antiretroviral therapy (cART) effectively suppresses HIV-1 replication but fails to achieve a cure due to the persistence of long-lived, resting memory CD4+ T cells that harbor latent and replication-competent HIV-1 DNA^1,2^. One potential curative strategy, known as “shock and kill,”^3^ involves the selective reactivation of HIV-1 gene expression using latency-reversing agents (LRAs), followed by the induction of cell death through virus-induced cytolysis or immune-mediated clearance of reactivated reservoirs^4–6^. Despite significant increases in plasma and cell-associated viral RNA following LRA treatment, clinical interventions have not yet resulted in substantial reductions in the latent reservoir^7–9^. Notably, some LRAs may inadvertently promote cell survival, counteracting attempts to eliminate infected cells^10^. This phenomenon might be attributed to compromised activation of intrinsic cell death pathways, leading to inefficient or no elimination of cells expressing viral gene products.

Upon reactivation of persistent HIV-1 infection by LRAs, viral proteins can interfere with apoptosis pathways, yielding varied outcomes, including activation, inhibition, or delay of cell death. For instance, the viral proteins Nef, Tat, and Vpr can impair apoptotic processes to promote cell survival and facilitate viral replication during the early stages of the viral life cycle. Conversely, in the late stages, the HIV envelope and Vpu proteins can promote apoptotic cell death^11^. Consequently, the balance between anti-apoptotic and pro-apoptotic cellular and viral proteins determines the fate of infected cells, influencing their survival or apoptosis^12^. Apoptosis is intrinsically regulated by members of the Bcl-2 family of proteins, which modulate apoptosis at the mitochondrial level. Key anti-apoptotic molecules in this family include Bcl-2, BcL-2 homologs Bcl-xL, Bcl-W, A1, and Mcl-1^13^. Inhibition of these proteins can lead to mitochondrial outer membrane permeabilization, resulting in cytochrome c release into the cytosol and subsequent activation of the caspase cascade that executes apoptosis^12,14^. Evidence suggests that increased expression of the anti-apoptotic protein Bcl-2 can protect latently infected cells from apoptosis^15^. Thus, pharmacological approaches aimed at triggering intrinsic apoptotic pathways in reactivated HIV-infected reservoir cells present a promising strategy for viral reservoir elimination^16,17^.

Pro-apoptotic compounds targeting Bcl-2 family proteins, extensively developed in cancer research^18^, offer potential repurposing opportunities for HIV-1 reservoir elimination. For example, Navitoclax (ABT263), a dual inhibitor of Bcl-2 and Bcl-xL, and Venetoclax (ABT199), a selective Bcl-2 inhibitor, were initially designed as small molecule inhibitors targeting Bcl-2 family proteins in cancer therapeutics and have been investigated for their ability to induce cell death in HIV-infected cells^19–21^. However, the utility of ABT263 in clinical settings is limited by severe side effects, including on-target, dose-limiting thrombocytopenia^22,23^. As the only FDA-approved drug that targets BCL-2 family protein^24^ and currently, under clinical trials (NCT04886622), Venetoclax has demonstrated limited efficacy in reducing viral reservoirs in ex vivo experiments when administered as a single agent^20^.

To mitigate the severe in vivo side effects associated with ABT263, Dr. Zhou’s group has developed a series of proteolysis-targeting chimeras (PROTACs)^25^ exhibiting enhanced anti-leukemia efficacy while minimizing off-target toxicity^26–28^. PROTACs are bifunctional small-molecule degraders comprising two active ligands connected by a chemical linker. One ligand is designed to selectively bind to a protein of interest, while the other engages an E3 ubiquitin ligase. The recruitment of the E3 ligase to the target protein facilitates the formation of a ternary complex, which leads to ubiquitination and subsequent degradation of the target protein by cellular proteasomes^25,29^. By selecting cell or tissue-specific E3 ligases, PROTACs represent an innovative technology in precision medicine, enabling the targeted degradation of proteins in cancerous or infected cells with alternative protein expression^29,30^. Leveraging the characteristic low expression of Von Hippel–Lindau (VHL) E3 ligase in human platelets^27^, Dr. Zhou’s group attached a ligand for the VHL E3 ligase to two different solvent-exposed rings of ABT263, leading to the discovery of two PROTACs, DT2216, and PZ703b, which exhibit selective cytotoxicity toward cancer cells while sparing platelets, thereby preventing thrombocytopenia^27,28^. Notably, both DT2216 and PZ703b have been shown to specifically degrade Bcl-xL, with concurrent Bcl-2 inhibitory activity, in lymphoblast cell lines derived from patients with lymphoblastic leukemia^27,28^. As DT2216 has recently entered clinical trials for the treatment of Bcl-xL-dependent relapsed or refractory peripheral T-cell lymphoma and cutaneous T-cell lymphoma (NCT04886622), the potential effects of DT2216 and PZ703b on the reduction of HIV-1 reservoirs remain to be explored.

Utilizing the advantages of protein degraders over traditional occupancy-based inhibitors allows for the circumvention of drug resistance arising from target protein overexpression or mutations, as protein degraders can degrade the entire protein^31,32^. Thus, this strategy has emerged as a promising antiviral approach to combat infectious diseases^33–36^, although it remains underexplored in the context of HIV-1 infection. In this study, we aim to repurpose currently developed Bcl-2/Bcl-xL PROTACs to investigate the potential of protein-degrading modalities to target cellular proteins for degradation, ultimately achieving host-directed antiviral effects.

## Results

### Evaluation of BCL-2/BCL-XL Antagonists for Reactivating and Selective Killing of HIV from Latency

To systematically evaluate the effectiveness of newly developed BCL-2/BCL-XL PROTACs and their predecessor small molecules in reactivating HIV from latency, we utilized J-Lat 10.6 cells, a well-established latently HIV-infected T cell line that expresses green fluorescent protein (GFP) upon HIV reactivation^37^. The expression of GFP correlates with viral production in response to latency-reversing agents (LRAs). J-Lat 10.6 cells and their parental Jurkat cells were treated with BCL-2/BCL-xL antagonists at varying concentrations for 48 hours. Dimethyl sulfoxide (DMSO) was used as the vehicle control, while TNF and Prostratin were used as positive controls for the treatment. The parental Jurkat cells provided a background control. The level of GFP expression upon HIV reactivation was measured using flow cytometry, reporting the percentage of cells exhibiting increased fluorescence relative to the total cell count (**Fig. 1a**). Compared to the approximately 55% HIV latency reversal observed in J-Lat 10.6 cells treated with TNF or Prostratin (**Fig. 1a, right**), PZ703b induced a maximum of 25% reactivation at the highest concentration, with effects diminishing at lower doses (**Fig. 1a, left, red**). Treatment with ABT263 resulted in detectable GFP induction, although at lower levels than PZ703b (**Fig. 1a, left, blue**). In contrast, DT2216 and ABT199 did not induce significant GFP expression throughout the treatment period (**Fig. 1a, left, orange and green**). Obatoclax, a known BCL-2 antagonist, has previously demonstrated the ability to reactivate latent HIV-1 in J-Lat 10.6 cells at micromolar concentrations^38^. However, fluorescent signals were also detected in parental Jurakt cells treated with Obatoclax, indicating spontaneous emission or excitation of signals (data not shown), rendering it unsuitable for this model. Notably, PZ703b and ABT263, which act as dual antagonists of Bcl-2 and Bcl-xL, achieved significant HIV reactivation at nanomolar concentrations, demonstrating superior potency compared to other screened BCL-2/BCL-xL antagonists.

**Fig. 1.**
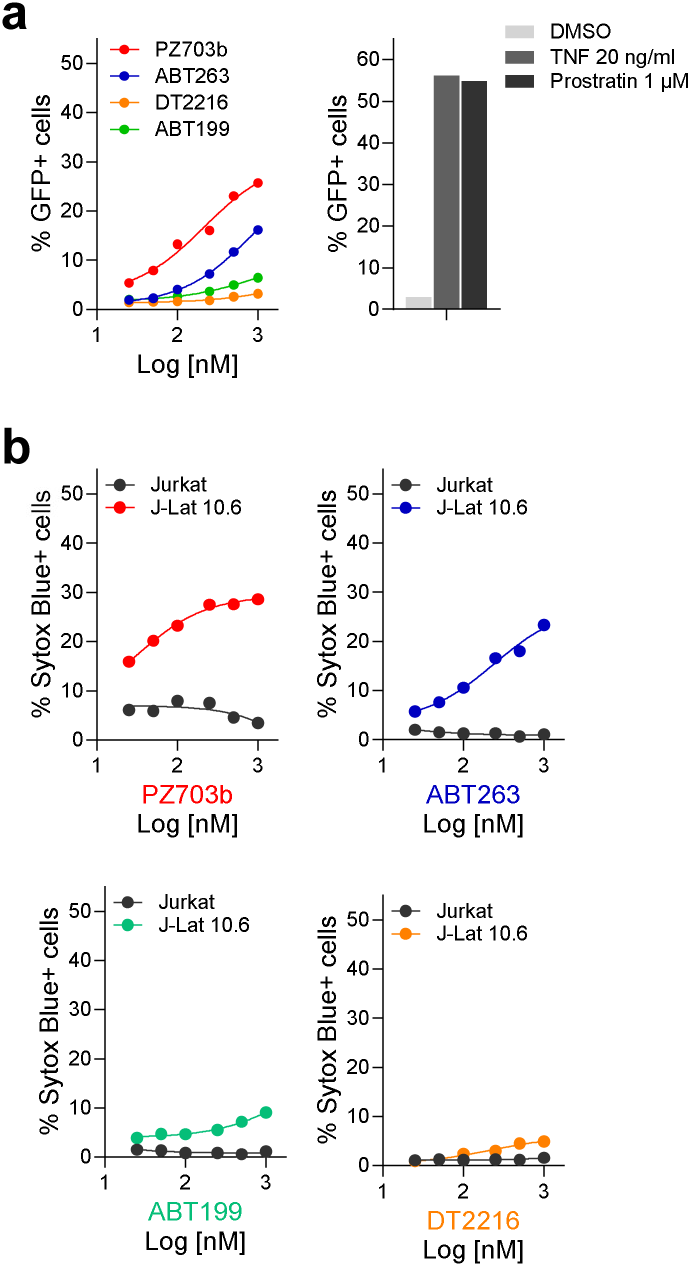
PZ703b Shows Potency in HIV-1 Reactivation and Preferential Killing of J-Lat Cells. Latently infected J-Lat 10.6 cells and their parental Jurkat cells were treated with serial concentrations of BCL-2/BCL-XL antagonists (PZ703b, ABT-263, ABT-199, and DT2216) for 48 hours. TNF and Prostratin were used as positive controls, while DMSO served as the negative control. **a** Reactivation of HIV-1 was quantified using flow cytometry, represented as the percentage of GFP-positive cells. **b** Cell death was assessed using SYTOX Blue, an impermeable nucleic acid dye that indicates compromised cell membranes. The percentage of SYTOX Blue-positive cells post-treatment was quantified by flow cytometry.

To further investigate whether PZ703b and the other antagonists induce cytopathic effects (CPE) in reactivated cells or cause cytotoxicity in all treated cells, we assessed cell death following the reversal of latent infection. The J-Lat 10.6 cell line contains a full-length integrated HIV genome, with the Nef gene replaced by the GFP open reading frame and the Env gene suppressed by a frameshift mutation. This configuration permits the production of viral proteins associated with CPE^37^. Both parental Jurkat and J-Lat 10.6 cells were treated and monitored for cell death using SYTOX Blue, a dead cell stain that penetrates and intercalates nucleic acids only when the cell membrane is compromised. Neither TNF nor Prostratin treatment resulted in a substantial fraction of SYTOX Blue-positive cells in either cell line, reflecting their latency-reversing capabilities without significant cytotoxic effects (data not shown). However, both PZ703b and its predecessor, ABT263, exhibited significantly higher toxicity to J-Lat 10.6 cells when treated for 48 hours (**Fig. 1b, upper two panels**), indicating that both compounds are highly effective in killing HIV-1 latently infected cells. DT2216 and ABT199 demonstrated slight cytotoxicity in J-Lat 10.6 cells compared to Jurkat cells (**Fig. 1b, lower two panels**). While well-known compounds such as Prostratin and Bryostatin can reverse HIV-1 latency without inducing significant CPE or cell death, additional strategies will be needed to eliminate reactivated latently infected cells. The results presented in Fig. 1 demonstrate that PZ703b is the first compound shown to effectively reverse latent infection and reduce the latent reservoir alone.

### PZ703b Reactivates Latent HIV Through the Noncanonical NF-kB Signaling Pathway

Our initial screening results indicate that dual antagonists of Bcl-2 and Bcl-xL can function as latency reversal agents (LRAs). To confirm the potency and efficacy of PZ703b and ABT263 in reactivating HIV-1 latency, we conducted experiments to measure the expression of HIV-1 transcripts and the HIV core protein Gag. We utilized real-time TaqMan quantitative PCR to assess HIV-1 transcript levels and flow cytometry to analyze HIV core protein expression before and after treatment with PZ703b or ABT263, with DMSO as a negative control and TNF as a positive control. In **Fig. 2a**, we evaluated whether PZ703b can activate latent HIV gene expression, suggesting full reactivation of HIV and subsequent viral production. After administering PZ703b for 24 and 48 hours, we observed a dose-dependent activation of HIV gag RNA transcription and latent HIV proviral RNA in J-Lat cells, compared to the negative control (DMSO) and the positive control (TNF) (**Fig. 2a, and Supplementary Fig. 1a**). Additionally, both PZ703b and ABT263 significantly increased the frequency of cells expressing the HIV core protein Gag and the reporter GFP in a dose-dependent manner (**Fig. 2b,c and Supplementary Fig. 1b**). Western blot analysis further confirmed the induced expression of HIV Gag p24 in J-Lat cells following treatment with higher doses of PZ703b (**Fig. 2d**). Collectively, these results demonstrate that PZ703b effectively activates the transcription of the latent HIV provirus in J-Lat cells, leading to the expression of both early gene products, such as Nef, and late gene products, such as Gag.

**Fig. 2.**
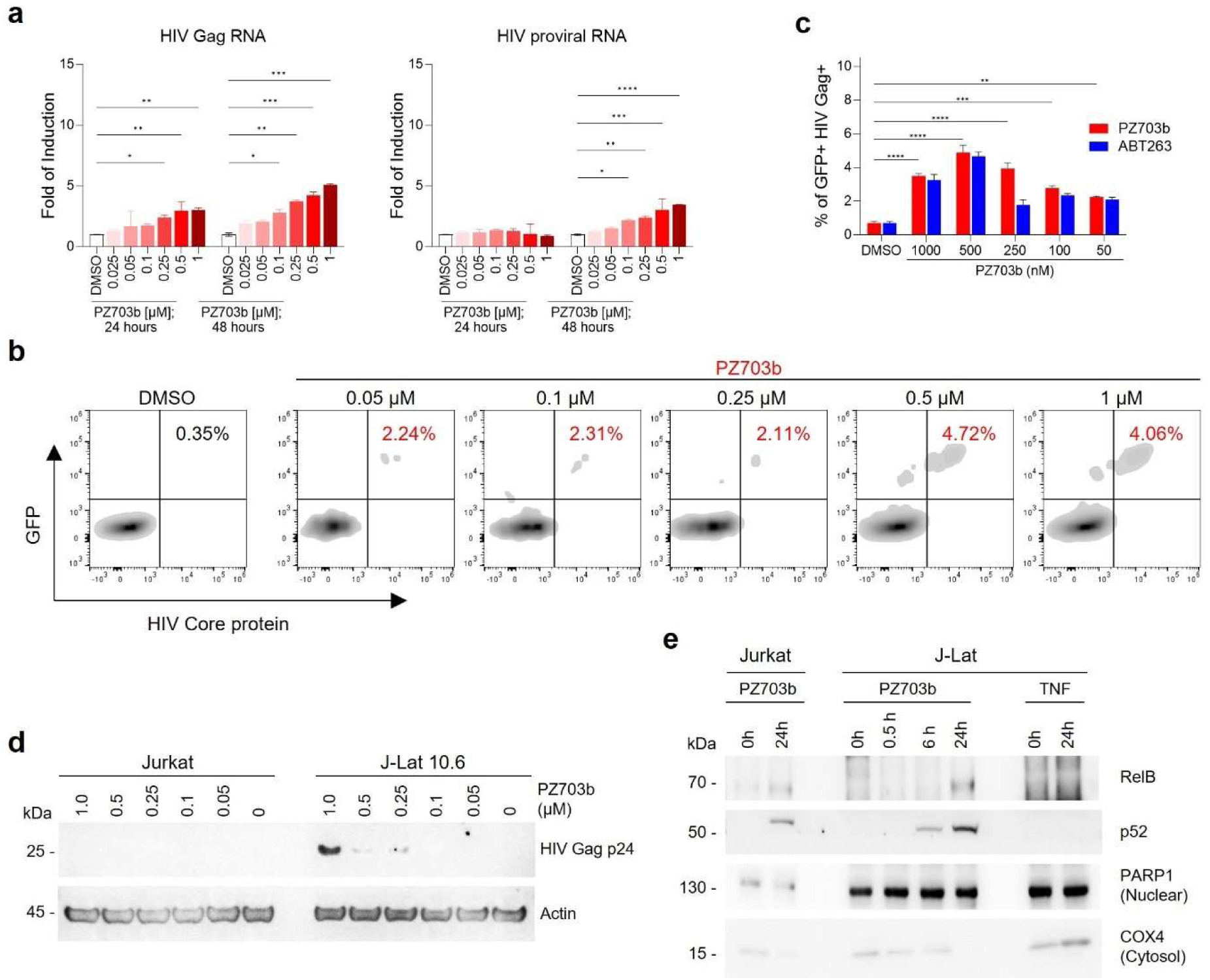
PZ703b reactivates HIV-1 latency in J-Lat cells. **a**. RNA isolated from cells treated as indicated was analyzed for HIV Gag and proviral RNA content. **b**. Intracellular staining of HIV-1 Gag proteins demonstrates the effectiveness of PZ703b in reversing latent viral infection. Cells were treated, harvested, fixed, and permeabilized for intracellular staining of HIV-1 Gag proteins using an RD1 (Phycoerythrin)-conjugated KC57 antibody. **c**. Data from panel **b** are duplicated and presented as mean ± SD, with P values ≤ 0.05 indicated by an arrow. **d**. Western blot analysis detected the expression of HIV Gag p24 protein upon increasing concentrations of PZ703b, with Actin serving as the internal control. **e**. Cells subjected to nuclear fractionation and treated as indicated were analyzed by western blotting. COX4 and PARP1 were used as controls for cytoplasmic and nuclear proteins, respectively.

Bcl-2 and Bcl-xL have previously been implicated in the regulation of NF-kB pathway^39,40^, which is one of the primary pathways targeted by a number of promising LRAs^41^. To investigate whether PZ703b activates the canonical NF-kB pathway, we treated J-Lat cells with DMSO or various concentrations of PZ703b for 24 hours, followed by intracellular staining for the phosphorylated p65 subunit of NF-kB and subsequent flow cytometry analysis. Notably, treatment with PZ703b resulted in a significant decrease in phosphorylated p65, which normally translocate to the nucleus to initiate HIV transcription, compared to the DMSO mock control and CD3/CD28- stimulated positive control (**Supplemental Fig. 1c**). These results suggest that PZ703b reactivates viral latency independently of the canonical NF-kB pathway. We next examined the role of the non-canonical NF-kB (NF-kB2) pathway, which functions as a slow, persistent, and stimulus-selective mechanism that may juxtapose canonical NF-kB signaling^42^ and could be responsible for the PZ703b-dependent transcriptional activation of HIV. We analyzed the nuclear translocation of NF-kB2 subunits following treatment with PZ703b, using PARP1 as a nuclear fraction marker and loading control, and COX4 as a cytosolic protein control (**Fig. 2e**). Our findings demonstrated that NF-kB2 subunits, p52 and RELB, translocated to the nucleus after 24 hours of PZ703b treatment. In contrast, TNF, known to activate the canonical NF-kB pathway, did not induce nuclear translocation of these subunits (**Fig. 2e**). These data suggest that PZ703b functions as an atypical latency-reversing agent (LRA), reactivating transcription of latent HIV proviruses through activation of the non-canonical NF-kB2 pathway, which may help minimize off-target effects^42,43^.

### Mechanisms of Selective Killing of HIV-Latently Infected Cells by PZ703b

PZ703b and its sibling compound, DT2216, are newly developed PROTACs (proteolysis-targeting chimeras) that demonstrate safer and more potent anti-cancer properties compared to their warhead, ABT263^27,28^. The target protein-binding moiety of PZ703b and DT-2216 is derived from ABT263^27,28^, which is a dual inhibitor of Bcl-2 and Bcl-xL. This rationale originally suggested that both PZ703b and DT2216 could target and degrade both Bcl-2 and Bcl-xL. However, studies have shown that PZ703b and DT2216 specifically degrade Bcl-xL but not Bcl-2 in lymphoblast cell lines and lymphomas^27,28,44^. This differentiation may be attributed to their lysine preference and the stability of the ternary complex formed for target protein ubiquitination and degradation^45^. In this study, we investigated the degradation efficiency and specificity of PZ703b and DT2216 on Bcl-xL and Bcl-2 protein levels in Jurkat and J-Lat cell lines. DT-2216 induced only Bcl-xL protein degradation with a similar DC_50_ (half-maximal degradation concentration) to prior findings^27^, without affecting Bcl-2 levels in either cell line (**Supplementary Fig. 2a,b**). In contrast, PZ703b effectively induced degradation of both Bcl-xL and Bcl-2 proteins after 24 hours of treatment, even at a concentration of 50 nM, demonstrating greater efficacy compared to DT2216 in both cell lines (**Fig. 3a**). The DC_50_ values for inducing degradation of Bcl-xL and Bcl-2 were approximately 10 nM and 50 nM, respectively, across both cell lines (**Fig. 3b**). This discrepancy is supported by findings that DT2216 can only form a stable complex with Bcl-xL, while PZ703b can bind both proteins, as shown by NanoBRET assays^27,28^. Our results highlight the significant efficacy and specificity of PZ703b in degrading Bcl-xL and Bcl-2 in Jurkat T lymphocytes. Although Jurkat cells are often used in leukemia studies, PZ703b’s efficacy and specificity in reducing Bcl-2 and Bcl-xL levels may vary by cell type and remain independent of HIV infection.

**Fig. 3.**
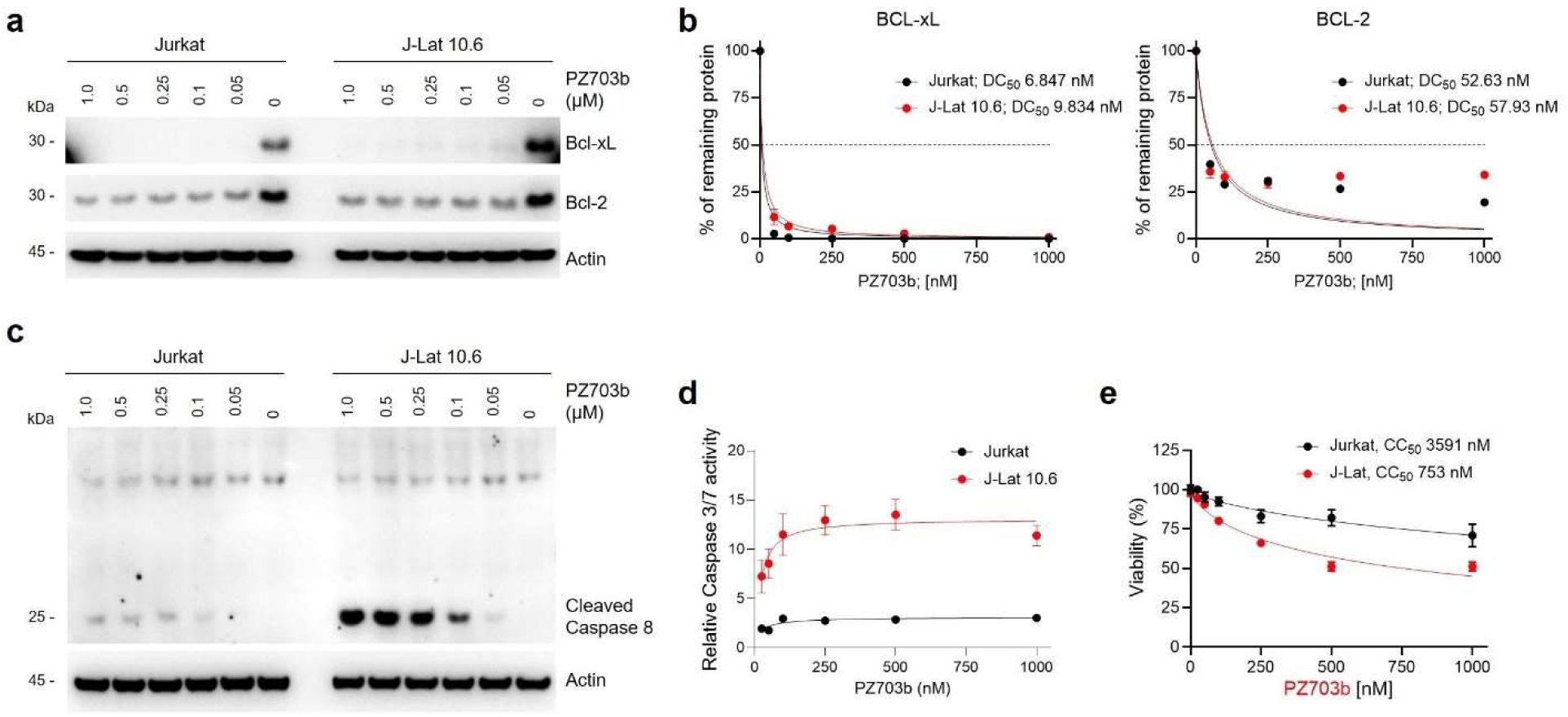
PZ703b degrades Bcl-xL and Bcl-2 proteins and selectively induces apoptosis in HIV-latently infected cells. **a**, **b** Degradation of Bcl-xL and Bcl-2 in Jurkat and J-Lat cells following treatment with PZ703b. Western blot analysis was performed to assess the protein levels of Bcl-xL, Bcl-2, and Actin after a 24-hour treatment with the indicated concentrations of PZ703b. DC50 indicates the drug concentration required for 50% protein degradation. Actin served as a loading control, with untreated sample densities normalized to 100% for comparison. Treated samples were referenced to Actin and then normalized to calculate the reduction percentage. **c** Activation of the caspase cascade, indicated by the presence of cleaved Caspase 8. Cells were treated as described in **a**, and lysates were analyzed using Western blotting with an anti-Caspase 8 antibody. The band size of 25 kDa corresponds to cleaved Caspase 8. **d** Enhancement of Caspase 3/7 activity in PZ703b-treated J-Lat cells. Jurkat or J-Lat cells were treated with increasing concentrations of PZ703b for 24 hours. Following treatment, cells were lysed and incubated with a caspase substrate (Promega). Luminescence was measured using a TECAN instrument after the incubation period. Two replicates were performed for each treatment group. **e** MTS assay in Jurkat and J-Lat cells, incubated with increasing concentrations of PZ703b for 24 hours. Data are presented as mean ± s.d. from three replicate cultures in a representative experiment.

Apoptosis is a programmed cell death process regulated by Bcl-2 and Caspase family proteins. Loss of Bcl-2 and Bcl-xL leads to mitochondrial outer membrane permeabilization and the subsequent release of cytochrome c into the cytosol, which initiates the self-cleavage of initiator Caspase 8 and activates the intrinsic apoptotic pathway. PZ703b, a novel dual degrader of Bcl-2 and Bcl-xL in Jurkat cells, downregulates both proteins, thereby inducing apoptosis. To assess the impact of PZ703b on intrinsic apoptosis activation, we measured the levels of active cleaved caspase 8 in PZ703b-treated Jurkat and J-Lat cells using western blotting. As shown in **Fig. 3c**, PZ703b significantly increased the levels of cleaved caspase 8 in J-Lat cells compared to Jurkat cells. Active Caspase 8 subsequently cleaves executor caspases, caspase 3 and 7, into their active forms. Therefore, we further evaluated whether PZ703b selectively enhances the activity of these executor caspases in latently HIV-1 infected cell lines. Jurkat and J-Lat cells treated with PZ703b were lysed, and the activities of Caspases 3 and 7 were measured by incubating the lysates with a substrate containing the tetrapeptide DEVD. Luminescence from PZ703b-treated cell lyastes was normalized against that of DMSO-treated control. As a result of the selective activation of Caspase 8 (**Fig. 3c**), PZ703b treatment led to a five-fold increase in the activation of Caspases 3 and 7 in J-Lat cells compared to Jurkat cells (**Fig. 3d**). These selective effects of PZ703b on J-Lat cells illustrate that PZ703b-mediated killing of latently infected T cells is linked to enhanced caspase cascade activation in an HIV-dependent manner, counteracting anti-apoptotic molecules to facilitate apoptotic pathways. We also conducted MTS and trypan blue staining assays to evaluate the selective apoptotic effects on cell viability. PZ703b demonstrated approximately five-fold greater cytotoxicity against J-Lat cells (half-maximal cytotoxic concentration (CC_50_) of 0.75 µM) compared to Jurkat cells (CC_50_ of 3.5 µM) in both the MTS assay (**Fig. 3e**) and the trypan blue exclusion assay (**Supplementary Fig. 2c**). In contrast, DT-2216 had no effect on the viability of either cell line in both assays (**Supplementary Fig. 2d**), indicating that selective killing activity requires the simultaneous reduction of Bcl-2 and Bcl-xL proteins in HIV-infected cells. Overall, PZ703b was established as a novel dual degrader of Bcl-2 and Bcl-xL, effective at selectively killing HIV-latently infected cells.

### PZ703b Does Not Lead to Global Activation of Primary T Cells

A critical criterion for evaluating promising therapeutic candidates for preclinical advancement in HIV reservoir treatment is their ability to avoid global T-cell activation, which can lead to cytokine release and toxicity. In this study, we investigated the impact of PZ703b treatment on T cell activation. Isolated human peripheral blood mononuclear cells (PBMCs) from two healthy donors were treated with PZ703b at varying concentrations for 72 hours. Control treatments included Prostratin, a protein kinase C (PKC) agonist, is known to trigger the activation of uninfected bystander T cells. This activation represents a significant disadvantage of current latency reversal agents (LRAs), as it may lead to unwanted immune activation and potential collateral damage to healthy cells. We assessed the surface expression of activation markers CD69, CD25, and HLA-DR on CD3-positive lymphocytes, including CD4+ and CD8+ T cells, from the treated PBMCs. The results shown in **Fig. 4** indicate that the expression of activation markers did not significantly increase following exposure to varying concentrations of PZ703b compared to the DMSO negative control. Importantly, PZ703b did not induce activation marker levels comparable to those observed with the positive control, Prostratin, which significantly increased activation markers during the study period. The results were consistent across both donors.

**Fig. 4.**
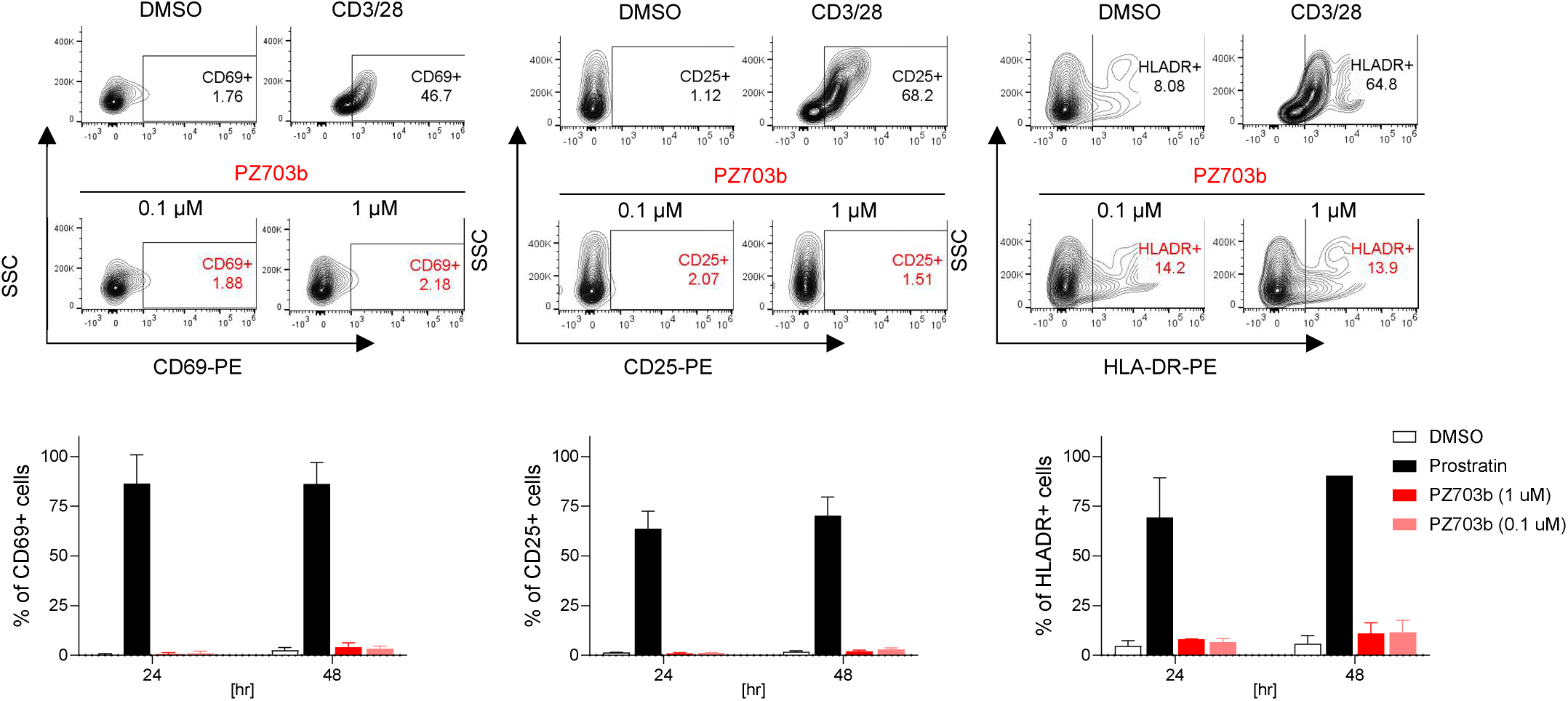
PZ703b does not lead to global activation of T cells. Peripheral blood mononuclear cells (PBMCs) obtained from healthy donors were treated for three days with 0.1% DMSO as a negative control, Prostratin as a positive control, or PZ703b. Flow cytometry was used to quantify the expression of activation markers (CD69, CD25, HLA-DR) on CD3+ T cells. The values represent the percentage of CD3+ cells expressing each individual activation marker, indicating the level of T cell activation.

### PZ703b Reverses Latency, Selectively Kills Reactivating Cells and Reduces Reservoir Size in the T_CM_ model of latency

In the HIV-1 life cycle, the expression of viral proteins can induce cytopathic effects, leading to cell death. For instance, the HIV envelope protein reduces BCL-2 levels, activating Bax and Bak, which form pores in the mitochondrial membrane, ultimately resulting in cytochrome C release and the activation of caspase cascades that drive apoptotic cell death^46,47^. Conversely, HIV protease-generated Casp8p41 induces apoptosis, which can be prevented by BCL-2^48,49^. Additionally, the HIV-1 accessory protein Vpu has been reported to suppress BCL-xL, significantly upregulated in individuals living with HIV^50^, to promote apoptosis^51^. These findings support our hypothesis that reactivating latently infected HIV-1 cells may synergize with PZ703b in promoting cell death to reduce the size of HIV reservoirs. To test this hypothesis, we conducted a central memory T cell (T_CM_) model of latency as described by Bosque’s group^52,53^. This model can mimic the environment of peripheral blood mononuclear cells (PBMCs) isolated from HIV-1 patients undergoing various ART regimens. Naive CD4+ T cells were isolated from HIV-negative donor peripheral blood mononuclear cells (PBMCs) via negative selection. The isolated CD4+ T cells were then activated with anti-CD3/CD28 Dynabeads in the presence of anti-IL-4, anti-IL-12, and TGF-β before being infected with HIV-1 laboratory-adapted replication-competent viruses. Following a total of six days of infection, we washed the infected cells and cultured them in a medium containing a combination of HIV integrase inhibitor Raltegravir (RLV) and viral fusion inhibitor Enfuvirtide (ENF) to drive the latency stage. After four days of culture in the presence of these antiretrovirals, latently infected central memory T cells were positively isolated for further study. We employed a cross-clade ultrasensitive real-time TaqMan PCR-based assay^54^ to confirm HIV latency in the T_CM_ model of latency. Latently infected cells containing integrated HIV-1 DNA exhibited an average Cq value of 25, indicating approximately 300 to 3,000 copies of HIV-1 latent proviruses^54^ in both HIV-LAI and HIV-IIIB latently infected primary T cells, whereas uninfected central memory T cell were undetectable (**Supplementary Fig. 3a)**.

To prevent new rounds of infection while allowing the completion of the viral life cycle, we maintained the presence of the two previous ARTs to assess whether PZ703b could activate latent HIV-1 and reduce the population of infected cells. After treating latently infected T_CM_ cells with PZ703b for 72 hours, we measured the level of reactivation by intracellular staining of the HIV-1 Core protein with KC57 antibody, followed by flow cytometric analysis. DMSO was used as a negative control, while anti-CD3/CD28 Dynabeads served as a positive control for T cell receptor co-stimulation. The BCL-2 inhibitor ABT-199, known for reducing the T_CM_ cells of latency, was also included in our analysis^16^. The subset of HIV-1 Core protein-positive cells was gated from the total cell population to determine the effect of PZ703b on the reactivation of latently infected T_CM_ cells. We found that neither PZ703b nor ABT-199 reactivated HIV-1 Core expression in cells latently infected with HIV-LAI or HIV-IIIB (**Supplementary Fig. 4a**), as compared to DMSO negative and anti-CD3/CD28 positive controls. While ABT-199 resulted in a substantial toxicity to the central memory T cells^20^ **(Supplementary Fig. 4b)**, this effect appeared to be primarily in HIV Core protein-negative cells **(Supplementary Fig. 4c)**. These results suggest that PZ703b is unlikely to exhibit synergistic effects with viral factors when used in conjunction with integrase inhibitors and fusion inhibitors, and therefore may not effectively reverse latency or selectively kill infected cells.

In the HIV life cycle, the protease enzyme is crucial for the maturation of viral proteins^55,56^. Evidence suggests that fully processed viral proteins, such as Nef and Vpr, may protect virus-infected cells from apoptosis while simultaneously inducing the death of surrounding bystander cells^57,58^. Moreover, the activity of HIV protease within host cells can result in the non-specific cleavage of various host proteins, including those involved in apoptosis^59–62^. Given these functions, viral protease can manipulate or alter host mechanisms to enhance viral survival. Therefore, treatment with a protease inhibitor has the potential to preserve critical host mechanisms, thereby allowing PZ703b to exert its intended effects. To explore this hypothesis, we combined the HIV integrase inhibitor Raltegravir (RLV) with the viral protease inhibitor Darunavir (DRV) instead. This strategy aims to evaluate the specific effects of PZ703b on the viral reservoir without the influence of mature viral proteins (**Supplementary Fig. 3b**). PZ703b-mediated induction of HIV Core protein expression was observed in both HIV-LAI and HIV-IIIB latently infected T_CM_ cells after 72 hours of treatment compared to the DMSO control (**Fig. 5a**). Furthermore, PZ703b led to a significant increase in cell death in the HIV Core-positive population (**Fig. 5b**) compared to the HIV Core-negative population, indicating selective killing of infected cells. Treatment with ABT- 199 resulted in HIV Core protein levels that remained comparable to those observed in DMSO controls across all time points (**Fig. 5a**), supporting previous studies that demonstrate the lack of latency reversal activity associated with ABT-199 in HIV infection. Although ABT-199 treatment induced approximately 50% cell death in HIV Core-positive cells in HIV-LAI infected cells (**Fig. 5b**), this effect was not selective, as ABT-199 displayed cytotoxicity against the entire HIV-LAI infected population (**Supplementary Fig. 5a, middle**). This non-specific toxicity is consistent with previous reports indicating that ABT-199 adversely affects uninfected T_CM_ cells as well^20^. Notably, ABT-199 did not promote selective cell death in HIV-1 IIIB latently infected T_CM_ cells (**Fig. 5b**) and lacked the non-specific toxicity observed in HIV-IIIB infection (**Supplementary Fig. 5a, right**). Costimulation with CD3 and CD28 markedly enhanced CD69 surface expression, signifying robust T cell activation; however, neither PZ703b nor ABT-199 exhibited significant immunomodulatory effects on infected or uninfected T_CM_ cells (**Supplementary Fig. 5b**). PZ703b treatment was associated with a reduction in the anti-apoptotic protein BCL-xL in both infected and uninfected T_CM_ cells (**Supplementary Fig. 5c**), consistent with observations in the J-Lat cell line (**Fig. 3**). These findings suggest that PZ703b is able to promote apoptotic cell death in cells with active HIV gene expression, though this effect is dependent on viral protease inhibition.

**Fig. 5.**
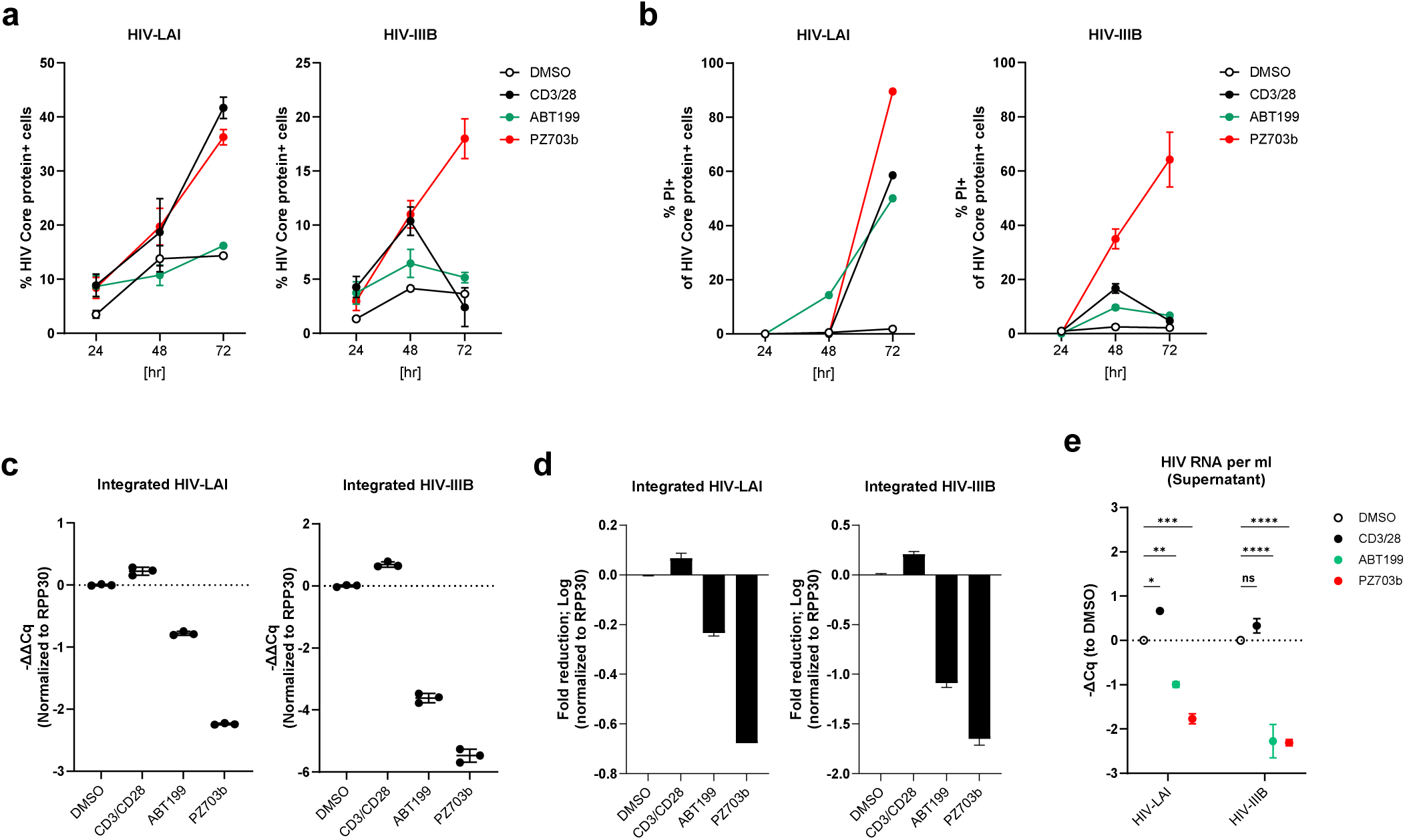
PZ703b reduces integrated HIV DNA in a T_CM_ model of latency through latency reversal and selective killing of reactivating infected cells. Naive CD4+ T cells were isolated from the peripheral blood mononuclear cells (PBMCs) of healthy donors and established in a latent state by infection with HIV-LAI or HIV-IIIB/H9, or left as uninfected controls, as detailed in the Methods section. **a** The percentage of cells expressing the HIV core protein was determined by intracellular staining with the KC57 antibody, followed by flow cytometry. **b** The percentage of apoptotic cell death among HIV core-expressing cells was assessed using propidium iodide staining, with apoptotic cells gated out from the HIV core-positive population. **c** Relative integrated HIV DNA from cells treated for three days was measured using nested TaqMan PCR. **d** The fold reduction in integrated HIV DNA after three days of treatment compared to the DMSO control was calculated and transformed using a logarithmic scale. **e** Relative viral production in the supernatant was assessed by measuring viral RNA release in three-day culture supernatants treated with indicated in the presence of antiretroviral therapies (ARTs), RAL and DRV. Cq values obtained from TaqMan RT-qPCR were normalized to DMSO. Data are presented as mean ± SDfrom two biological replicates. Statistical significance was determined using two-way ANOVA: *, p < 0.05; **, p < 0.01; ***, p < 0.001; ****, p < 0.0001.

To further demonstrate that PZ703b induces selective killing of latently infected cells, we extracted cell-associated DNAs to quantify integrated HIV-1 DNA, thereby assessing the impact of PZ703b on the reduction of the latent HIV-1 reservoir. Consistent with the observation that 60-80% of HIV Core-positive cells were apoptotic (**Fig. 5b**), PZ703b significantly decreased the amount of integrated HIV-1 DNA. This reduction was quantified as changes in ΔΔCq values, normalized to the internal gene control RPP30 and DMSO control, in both virus-infected cell types (**Fig. 5C**). Notably, this represented an approximate two-log decrease in the HIV-IIIB infected T_CM_ cells (**Fig. 5d**). In contrast, ABT-199 only modestly reduced the size of the infected T_CM_ cells (**Fig. 5c,d**) irrespective of HIV Core protein expression levels (**Fig. 5a**). Furthermore, CD3/CD28 costimulation resulted in a slight increase in integrated viral DNA compared to DMSO controls (**Fig. 5c,d**). Overall, these findings demonstrate that PZ703b has a more specific elimination effect in the T_CM_ model of latency.

To ascertain whether PZ703b influences the viral dissemination to surrounding uninfected T cells to HIV-1 infection, supernatants were collected for reverse transcriptase-PCR analysis to evaluate HIV RNA levels. While overall viral production was limited in the presence of ARTs, PZ703b further suppressed the release of viral particles, evidenced by reduced HIV RNA levels in the supernatant (**Fig. 5e**). This reduction in viral release aligns with the decreases in integrated HIV DNA observed (**Fig. 5c,d**), indicating that PZ703b possesses significant clinical potential for restricting the viral dissemination and reducing the size of the HIV reservoir *ex vivo*.

### Ex vivo treatment with PZ703b reduces HIV-1 reservoir

The persistence of resting CD4+ T cells harboring integrated HIV-1 genomes poses a significant challenge to cure HIV-1 due to their ability to produce a replication-competent virus. In ongoing HIV-cure-directed studies involving antiretroviral therapy (ART)-suppressed patients, the IPDA^®63^ has emerged as a promising strategy to measure replication-competent virus in reservoirs by distinguishing intact from defective proviruses, using a relatively small number of cells^64^.

Following our observations of PZ703b’s effectiveness against infected cells in the T_CM_ model of latency (**Fig. 5**), we aimed to evaluate its potency in a preclinical setting. We tested whether PZ703b could drive the reduction of latent HIV ex vivo in CD4+ T cells isolated from four HIV- positive female adults with HCV coinfection, all of whom were on ART (**Supplementary Table I**). The cells were treated with either DMSO or PZ703b in the presence of ART agents, including the integrase inhibitor RAL and the protease inhibitor DRV, for three days. After treatment, the cells were subjected to the IPDA assay to measure the frequency of intact provirus-positive cells. The IPDA quantifies intact HIV-1 proviruses as well as viral genomes with common defects, such as large internal deletions and hypermutation, which constitute the majority of reservoirs in people living with HIV^63,65,66^. While PZ703b may be capable of eliminating ex vivo reservoirs through the induction of apoptotic death in infected cells, we did not remove dying or dead cells by isolating Annexin V-negative CD4+ T cells prior to the IPDA assay^67,68^. Results indicated that the frequency of intact provirus-positive CD4+ T cells was significantly lower in the PZ703b-treated group compared to the DMSO control group (**Fig. 6a, left**). Three out of four participants experienced a reduction in intact viral copies per million cells, with an average reduction of 50% in intact provirus levels observed in the PZ703b group relative to DMSO treatment (**Fig. 6b**). Moreover, no significant changes were noted in the frequencies of CD4+ T cells harboring 5’- or 3’-defective proviruses among the four participants when comparing PZ703b treatment to DMSO (**Fig. 6a, middle and right**). Most defective proviruses have mutations in the tat and env genes, hindering full reactivation of viral gene expression and the production of viral proteins^65,66^, which may result in cells with defective HIV-1 being less capable of intrinsic apoptosis activation and again implicates the role of viral reactivation in PZ703b-mediated reservoir reduction. Thus, the observed decline in the frequency of intact provirus+ CD4+ T cells can be attributed to PZ703b- mediated cytotoxicity against HIV-1-infected cells carrying replication-competent viral genomes. The post-recovery cell viability of ex vivo repository samples tested was relatively modest (**Supplementary Fig. 6a**), resulting in an increased shearing index in both the DMSO and PZ703b groups (**Supplementary Fig. 6b**). However, we saw no discernable difference in either viability or DNA shearing between PZ703b treated cells and DMSO control treated cells, suggesting that the assay outcomes were not impacted. These findings provide direct evidence that ex vivo treatment with PZ703b eliminates latently infected CD4+ T cells of PWH on ART.

**Fig 6.**
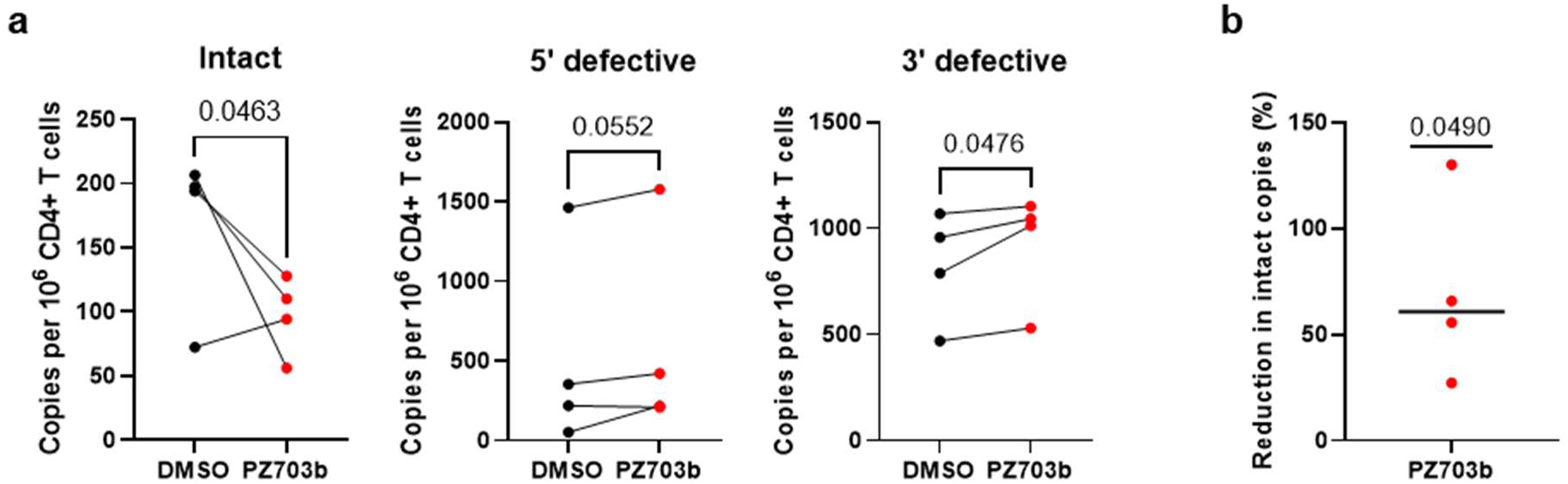
Ex vivo treatment with PZ703b reduces levels of intact HIV proviral DNA in samples from PWH on ART. **a** Frequencies of intact provirus+ CD4+ T cells (left), 5’-deleted provirus+ CD4+ T cells (middle), and 3’-deleted provirus+ CD4+ T cells (right) after treatment with PZ703b for three days. Data are reported as intact HIV-1 copies per million CD4+ T cells for each participant. Significance was determined by a one-tailed unpaired t-test. **b** Average frequency of intact provirus+ CD4+ T cells (top left) relative to DMSO. Data is reported as a percentage reduction (mean ± s.d.) from four participants. Significance was determined by one-sample t-tests.

The relationship between HCV coinfection and HIV-1 reservoir size in cART-treated HIV patients is complex and not fully understood. Some studies suggest that HCV coinfection may lead to an increased HIV-1 reservoir size, potentially due to chronic inflammation and immune activation^69^. However, other studies have found no association between HCV coinfection and HIV reservoir size^39^. Despite these mixed findings, ex vivo treatment with PZ703b has limited effects on the reduction of latent HIV in CD4+ T cells isolated from HCV-negative female adults (**Supplementary Fig. 6c**). Given that HCV co-infection appears to sensitize latently infected CD4+ T cells to apoptosis in untreated HIV-positive patients^70^, our preclinical observations underscore the critical need to further investigate the interactions between HIV and HCV infections and their influence on PZ703b-mediated ex vivo reduction of the HIV reservoir.

## Discussion

The “shock and kill” strategy aimed at eliminating the HIV-1 latent reservoir in CD4+ T cells rely on inducing latency reversal and viral reactivation, leading to either infected cell death or elimination by immune effector cells. In this study, we identified PZ703b, a novel protein degrader which like its parent compound ABT263, functions as a dual inhibitor of Bcl2 and Bcl-xL through protein degradation in Jurkat and J-Lat cell lines. We demonstrated that PZ703b and ABT263 effectively promote potent and specific caspase activation in a latently infected T cell line while sparing uninfected cells. Notably, ex vivo treatment with PZ703b in CD4+ T cells from people with HIV (PWH) eliminated up to 50% of cells containing intact proviruses, highlighting its potential as a promising therapeutic for reducing the HIV-1 reservoir. Given that ABT263 failed in phase II clinical trials due to on-target platelet toxicity, PZ703b offers a promising strategy for clinical development, demonstrating a favorable on-target profile with minimal platelet toxicity in the context of HIV reservoir reduction. Overall, our study positions PZ703b as the first-in-class protein degrader with potential applications in reducing reservoir size during chronic HIV infection.

To date, more than 160 compounds have been identified with latency-reversing activity; however, none of these latency-reversing agents (LRAs) have demonstrated a reduction in the size of the HIV reservoir or a delay in viral rebound in vivo^7,8,71^. Among these, pro-apoptotic molecules have been shown to activate NF-κB pathways and reactivate HIV-1 latency. SMAC mimetics, such as LCL-161, SBI-0637142, birinipant, and AZD5582, which target host anti-apoptotic factors like XIAP and cIAP1/BIRC2, serve as potent latency reversing agents (LRAs) by activating non-canonical NF-κB pathways^72,73^. This connection between apoptosis and latency reversal presents a promising strategy for reactivating the viral reservoir and promoting its selective elimination. In this study, PZ703b is established as the first pro-apoptotic protein degrader with demonstrated HIV reversal activity, specifically inducing selective toxicity towards HIV-infected cells and reducing the size of the ex vivo reservoir by 50%. Future studies could explore the potential synergistic effects of combining PZ703b with a SMAC mimetic.

In our study, we evaluated a range of pro-apoptotic molecules in the context of HIV infection, focusing on BCL-2/BCL-xL antagonists developed as cancer therapeutics and advanced to clinical trials. ABT-199 (Venetoclax), a selective BCL-2 inhibitor, has demonstrated efficacy in treating BCL-2-dependent hematological cancers^74^ and is FDA-approved^24,75^, However, our results indicated that it only modestly reduced HIV-1 levels in latently infected J-Lat cell line and HIV-infected primary CD4+ T cells. Obatoclax (GX 15-070), a pan-BCL-2 family inhibitor, is capable of inducing apoptosis in cancer cells and reactivating HIV-1; however, we found that its effectiveness on latently infected cells varies depending on the methods employed^38^. In contrast, ABT-263, which inhibits both BCL-2 and BCL-xL, was found to be less potent than PZ703b in reversing viral latency and selectively killing J-Lat 10.6 cells. This underscores the necessity of simultaneously targeting both proteins for effective “kick-and-kill” strategies. PZ703b, a derivative of ABT-263, exhibited dual-targeting capability by effectively degrading both BCL-2 and BCL-xL proteins in Jurkat cells and Jurkat-derived HIV-latently infected cell lines. While PZ703b was confirmed as a selective BCL-xL degrader in lymphoblast cell lines^28^, our findings indicate that differences between cell lines may influence its dual effects. Moreover, PZ703b exhibited enhanced activity at lower concentrations compared to ABT-263, highlighting the advantages of PROTACs over traditional small molecules^21^. It would be intriguing to further investigate the successors of PZ703b, specifically 753b and WH244, to evaluate their effectiveness in the same experimental framework as this study^26,45^. Overall, PZ703b presents a compelling therapeutic candidate for curative strategies against HIV-1 due to its specific targeting of HIV-infected cell death.

The study by Shan et al. illustrated the disconnect between results obtained from primary-cell models and clinical efficacy, prompting researchers to focus on ex vivo CD4+ T cells from ART- treated donors^10^. In our study, we tested the ex vivo potency of PZ703b and demonstrated that it effectively reduces the ex vivo HIV reservoir from HIV/HCV coinfection patients, achieving nearly a 50% elimination of reservoir cells within three days of treatment. This result indicates that PZ703b’s efficacy is consistent across both in vitro cell lines and ex vivo reservoirs. Current studies indicate that while individual latency-reversing agents (LRAs) demonstrate significant activity in vitro, they lead to only minimal increases in HIV-1 RNA transcripts in ex vivo patient samples^68,76^. This modest reactivation may result in a partial decrease in the viral reservoir but does not achieve complete elimination. To further explore the ex vivo efficacy of PZ703b, future work will investigate its ability to reverse HIV latency in ex vivo settings, comparing its effects to those of PMA/I-treated cells, which are the standard for evaluating maximum reactivation^68,77–79^. Overall, this study establishes PZ703b as the first protein degrader identified as a latency-reversing agent (LRA), demonstrating potent efficacy in promoting HIV latency reversal and selectively inducing death in infected cells as a monotherapy.

## Method

### Cell line culture

Jurkat and J-Lat full-length clone 10.6 cell lines were obtained through the NIH AIDS Research and Reference Reagent Program ^37^ and maintained in RPMI-1640 GlutaMax supplemental with 10% of heat-inactivated fetal bovine serum and 1x MycoZap (Roche).

### Human cell isolation and culture

Primary peripheral blood mononuclear cells (PBMCs) were acquired from the New York Blood Center in Long Island City from de-identified healthy donors. CD4+ T cells were purified using the EasySep Negative selection kit (STEMCELL). Cells were then cultured in RPMI-1640 GlutaMax supplemental with 10% heat-inactivated fetal bovine serum and 1x MycoZap (Roche) without any cytokines or stimuli. Cryopreserved PBMCs were obtained from participants in a cohort study of postmenopausal women with HIV led by Dr. Yin at Columbia University (IRB AAAF1644 and AAA2194^80,81^). Selection criteria included undetectable plasma HIV-1 RNA levels (<50 copies per ml). In total, samples from four women with HIV and Hepatitis C (HCV) coinfection were utilized in this analysis (see Supplementary Table 1). This protocol was approved by the Columbia University institutional review board. All participants provided written informed consent. Leukapheresis samples from healthy HIV− participants were obtained from the New York Blood Center in Long Island City. PBMCs were purified from leukapheresis samples by density gradient centrifugation.

### In vitro cell line model for screening BCL-2/BCL-XL antagonists

Jurkat or J-Lat 10.6 cells were seeded in 24-well plates, with 1 million cells per milliliter per well. The cells were treated with either TNF (20 ng/ml) or Prostratin (1 µM) as negative and positive controls, respectively. Alternatively, they were treated with BCL-2/BCL-XL antagonists at serially diluted concentrations or with diluent control (DMSO). After 24, 48, or 72 hours of treatment, the cells were harvested for GFP reporter measurement and assays.

### Measurement of supernatant HIV-RNA by TaqMan PCR

Viral particles were centrifuged at 20,000g for 2 hours at 4°C to pellet. The viral pellet was resuspended in 100 µL of residual medium, and viral RNAs were extracted using the QIAamp Viral RNA Kit (Qiagen) following the manufacturer’s instructions. RNA concentrations were quantified, and 500 ng of viral RNA were mixed with 0.5 µM forward primer, 0.5 µM reverse primer, and 0.2 µM probe for qPCR, using the TaqMan™ Fast Virus 1-Step Master Mix (Thermo). Samples were analyzed on a QuantiStudio 3 real-time TaqMan PCR system with default cycling parameters. Each run included tenfold serial dilutions of an RNA standard and negative control wells.

### Protein expression analysis using immunoblotting

As indicated in the figures, cells were treated with 0.1% DMSO (Sigma) or serially diluted PZ703b (medchemexpress) for 24 hours. After treatment, cells were lysed and subjected to Western blot analysis. Cell lysates (20 to 50 µg) were run on 4-12% NuPAGE Bis-Tris protein gels (Invitrogen) and transferred to a polyvinylidene difluoride (PVDF) membrane (BioRad) overnight at 30 volts in NuPAGE™ Transfer Buffer containing 10% ethanol and 0.05% SDS. The membranes were blocked in PBS (phosphate-buffered saline) with 5% bovine serum albumin (Sigma) for one hour at room temperature. Following blocking, membranes were incubated with primary antibodies at appropriate titers, as specified in the data sheets, overnight at 4°C. After washing three times with PBST (PBS with 0.5% Tween 20), membranes were hybridized with horseradish peroxidase (HRP)-conjugated secondary antibodies for two hours at room temperature. After three additional washes with PBST, membranes were treated with Super Signal West Pico PLUS Chemiluminescent Substrate (Thermo), and chemiluminescence was detected using a BioRad ChemDoc system. Antibodies used for immunoblotting were mouse anti-human Caspase 8 antibody (Cell Signaling, #5668), rabbit anti-human BCL-2 monoclonal antibody (Cell Signaling, #2118), rabbit anti-human BCL-xL monoclonal antibody (Cell Signaling, #7864), rabbit anti-Actin monoclonal antibody (Cell Signaling, #2859), NF-κB2 p100/p52 antibody (Cell Signaling, #4882), rabbit anti-human RelB (C1E4) monoclonal Ab (Cell Signaling #4922), monoclonal anti-HIV p24 antibody (NIH AIDS Reagent Program, #530), mouse anti-human PARP (Santa Cruz, #sc-8007), mouse anti-human COX (Santa Cruz, #sc-376731).

### Cytotoxicity MTS Assay

Jurkat or J-Lat 10.6 cells were seeded in 24-well plates, with 1 million cells per milliliter per well. The cells were treated with DMSO as a negative control or BCL-2/BCL-XL antagonists at serially diluted concentrations. After 24, 48, or 72 hours of treatment, the cells were harvested for the assay. To evaluate the cytotoxicity effect of screened BCL-2/BCL-XL antagonists, a tetrazolium compound [3-(4,5-dimethylthiazol-2-yl)-5-(3-carboxymethoxyphenyl)-2-(4-sulfophenyl)-2H- tetrazolium, inner salt] (MTS) assay (Promega G3582) was used. Colorimetric MTS assay can determine cell proliferation, viability, and cytotoxicity. The quantity of formazan product, as measured by the amount of 490nm absorbance, is directly proportional to the number of living cells in the culture. The absorbance of each sample was read by a microplate reader (TECAN), using a Magellan software program.

### Measurement of Caspase activity

Jurkat or J-Lat 10.6 cells were seeded in 24-well plates, with 1 million cells per milliliter per well. The cells were then with BCL-2/BCL-XL antagonists at serially diluted concentrations, or with diluent control (DMSO). After treatment for 24, 48, or 72 hours, the cells were harvested for the assay. The Caspase 3/7 assay was performed by following the protocol provided by the manufacturer (Promega).

### Flow cytometry

After DMSO or PZ703b treatment for 24, 48, or 72 hours, cells were fixed, permeabilized, and stained intracellularly with antibodies according to Thermo intracellular staining protocol. The following antibodies were used for flow cytometry: phycoerythrin (PE)-conjugated mouse anti-HIV core protein antibody (Beckman Coulter, KC57-RD1), phycoerythrin (PE)-conjugated mouse anti-human BCL-xL protein antibody (Invitrogen), phycoerythrin (PE)-conjugated rabbit anti-human NF-kB p65 (Ser536) antibody (MACS, #12AD). GFP-positive cells were determined using DMSO-treated J-Lat cells as a negative control for downstream analysis. Cell deaths were measured using Live/Dead Fixable Violet dead cell stain (Invitrogen) or Annexin V/PI (BD) kits according to the manufacturer’s protocol. Gating for drug-induced cell death was based on uninfected and DMSO-treated controls. Fluorescence-activated cell sorting (FACS) analysis was performed on Attune™ NxT Flow Cytometer (Invitrogen). FACS data were analyzed using FlowJo software (Tree Star).

### Generated and cultured central memory CD4+ primary T Cells

Central memory CD4+ primary T Cell model was generated as previously described ^52^. Peripheral Blood Mononuclear Cells (PBMCs) were purified from the blood of HIV-negative donors using density gradient centrifugation (STEMCELL). The Human Naïve T Cell Enrichment Kit (STEMCELL) was used to isolate resting CD4+ T Cells. The isolated resting CD4+ T Cells were then maintained in RPMI-1640 GlutMax medium, supplemented with 10% heat-inactivated FBS, 1x Pen/Strep, and 1x MyxoZap (Roche). To activate and expand the resting T Cells, a medium containing Dynabeads human T-activator CD3/CD28 (Invitrogen), 2mg/ml anti-human IL-12 (PeproTech), 1mg/ml anti-human IL-4 (PeproTech) and 10 ng/ml transforming growth factor b1 (TGF-b1) was used. After three days of activation, Dynabeads were removed, and the cells were further cultured in a medium supplemented with 30 IU/ml of IL-2 (R-10-30) for expansion. The medium was refreshed on days 4 and 5.

### Infection of central memory CD4+ primary T Cells to establish a model of latency

Infection of replication-competent HIV-1 virus to establish latency in central memory CD4+ primary T Cells followed the method described by Bosque and colleagues^53^. On day 7 of the central memory T Cells generation protocol mentioned above, some cells were intentionally infected with HIV-IIIB or -LAI viruses by spinoculating them with 150 TCID50 at 2,000xg for two hours at 37°C. After that, the infected cells were separated from the supernatant, suspended in a fresh medium containing IL-2, and returned to the incubator. Meanwhile, the remaining cells were cultured as an uninfected control. On the 10th day, the cells were counted, washed, and suspended in a medium containing IL-2 at 1 million cells/ml cell density to achieve crowd infection. On the 13th day, the cells were washed and suspended in a fresh medium containing IL-2, 1 µM of Raltegravir, and 0.5 µM of either Darunavir or Enfuvirtide to drive latency. On the 17th day, the CD4-positive cells were isolated through magnetic positive selection using the STEMCELL manufacturer’s instructions and were then ready for the assay.

### Analysis of activation markers in human primary T Cells

PBMCs were isolated from identified healthy donor blood. Human primary T Cells were purified from human PBMC following the manufacturers’ manual (STEMCELL). Surface expression of CD69, CD25, and HLA-DR activation markers were analyzed by flow cytometry. Cells were cultured for 24, 48, and 72 hours with the vehicle control DMSO or the following drugs: TNF, PZ703b, or ABT199 or costimulated with anti-CD3/CD28 antibody beads (Invitrogen). Each treatment contained one million cells. Post-treatment, fluorophore-conjugated antibodies targeting CD69, CD25, or CD69 were used to stain. Antibodies were incubated with cells at 4 degrees for 30 to 60 minutes, then washed with PBS at least twice before running flow cytometric analysis with ThermoFisher Attune NxT Flow Cytometer.

### Measurement of total cell-associated integrated HIV genome

DNA was extracted from at least one million cells using a Puergene DNA kit (Qiagen) according to the manufacturer’s protocol and quantified using a NanoDrop spectrophotometer. Levels of total and integrated HIV DNA were measured as previously described with slight modification. Briefly, a first-step PCR mixture containing two forward primers specific for the human Alu elements and a reverse primer specific for the HIV Gag gene was followed by a nested real-time TaqMan PCR with a pair of forward and reverse primers and a probe able to amplify both total and integrated HIV genome. TaqMan Quantitative PCR was performed using HIV primers and probers cross-clade detect for HIV integrated and total DNA and a primer-probe set for RPP30 (RNase P) as an internal control. PCR reaction mixtures were prepared by mixing 500 ng of purified DNA, 1000 nM primers, and 250 nM probe with TaqMan™ Gene Expression Master 2x Mix (Applied BioSystem). The PCR reaction was then conducted in a QuantiStudio 3 real-time PCR system with the default condition setting. The Cq values of HIV DNA were normalized to the Cq values of RPP30. Integrated or total HIV DNA was calculated and expressed in one million CD4+ T cells by the standard number of HIV DNA copies from serial dilution of ACH2 DNA standard controls.

### Ex vivo elimination of latent reservoir

CD4+ T cells were isolated from cryopreserved PBMCs of participants on suppressive ARTs (Supplementary Table 1) and plated in cultured in RPMI-1640 GlutaMax supplemental with 10% heat-inactivated fetal bovine serum and 1x MycoZap (Roche) with PZ703b and ARTs (10 M DRV and 10 M RAL) and incubated at 37 °C for 72 hours. The CD4+ T cells were then washed with complete RPMI medium to remove the treatment and subjected to IPDA assay.

### IPDA® assay

An in-depth description of the IPDA rationale and procedure is available in Bruner et al.^63^. In this study, the IPDA was performed by Accelevir Diagnostics under company standard operating procedures. Cryopreserved PBMCs from each participant were thawed, and total CD4^+^ T cells were obtained via immunomagnetic selection (EasySep Human CD4^+^ T cell Enrichment Kit, Stemcell Technologies), with cell count, viability, and purity assessed both before and after selection. An average of two million untouched CD4^+^ T cells were recovered for each sample and equally divided for DMSO or PZ703b treatment. Genomic DNA was isolated using the QIAamp DNA Mini Kit (Qiagen). DNA concentrations were determined by fluorometry (Qubit dsDNA BR Assay Kit, Thermo Fisher Scientific), and DNA quality was determined by ultraviolet-visible (UV/VIS) spectrophotometry (QIAxpert, Qiagen). Genomic DNA was then analyzed by IPDA^®^. Up to 700 ng of genomic DNA was analyzed for each Proviral Discrimination reaction, while Copy Reference/Shearing reactions used diluted (1:100) samples, with a maximum of 7 ng of genomic DNA analyzed per reaction. Samples were processed and analyzed in batches by blinded operators.

### Statistical methods

GraphPad PRISM version 10 software was used for statistical analyses, and the statistical analysis methods are reported in the figure legends. *P* values of ≤0.050 were considered significant.

## Author Contributions

L.C.C. designed and conducted the experiments, generated and analyzed the results, coordinated collaborative efforts, and drafted, edited, and finalized the manuscript for submission. M.T.Y. and J.S. were responsible for the collection of human specimens, provided patient histories for discussion, advised on handling human specimens, and contributed to manuscript editing. G.M.L. and K.D.R. worked on the IPDA assay of human ex vivo samples; G.M.L. also advised on the interpretation of IPDA data and helped revise the manuscript for submission. A.K.D. initiated the concept of repurposing Bcl-2/Bcl-xL PROTACs for HIV cure studies, discussed experimental results, facilitated collaboration among researchers, and contributed to manuscript editing for submission.

