## Supplemental Materials for "A First-in-Class Dual Degrader of Bcl-2/Bcl-xL Reverses HIV Latency and Eliminates Ex Vivo Reservoirs"

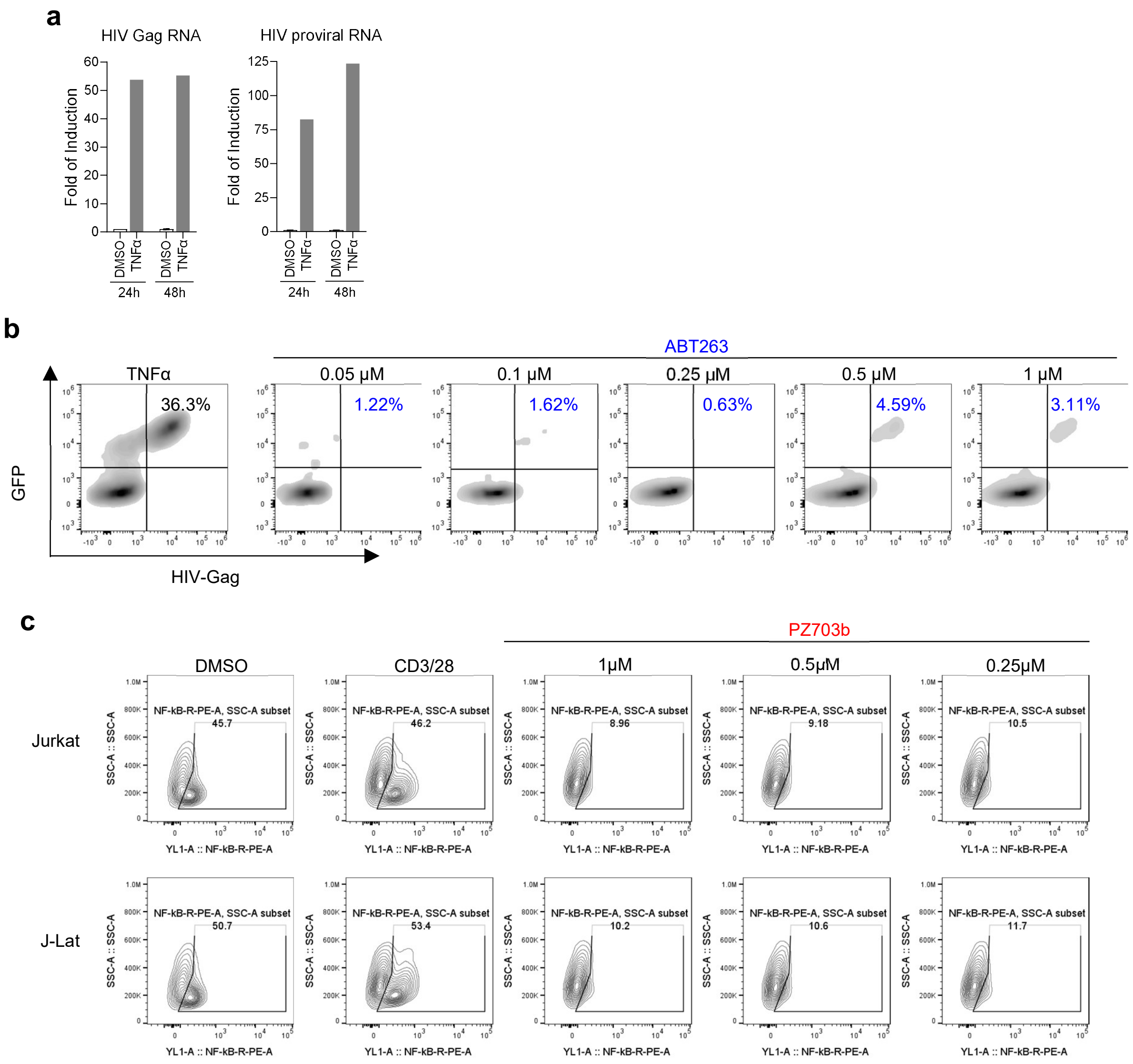


**Supplementary Fig. 1 ABT263 Reactivates HIV Latency in J-Lat Cells, and PZ703b Represses Canonical NF-kB Pathway Signals a** RNA isolated from cells treated as indicated was analyzed for HIV Gag and proviral RNA content. **b** Intracellular staining of HIV-1 Gag proteins demonstrates the effectiveness of PZ703b in reversing latent viral infection. Cells were treated, harvested, fixed, and permeabilized for intracellular staining of HIV-1 Gag proteins using an RD1 (Phycoerythrin)-conjugated KC57 antibody. **c** PZ703b treatment resulted in decreased NF-kB activity compared to DMSO and CD3/CD28 stimulation controls, which is consistent with its effects on T cell activation.

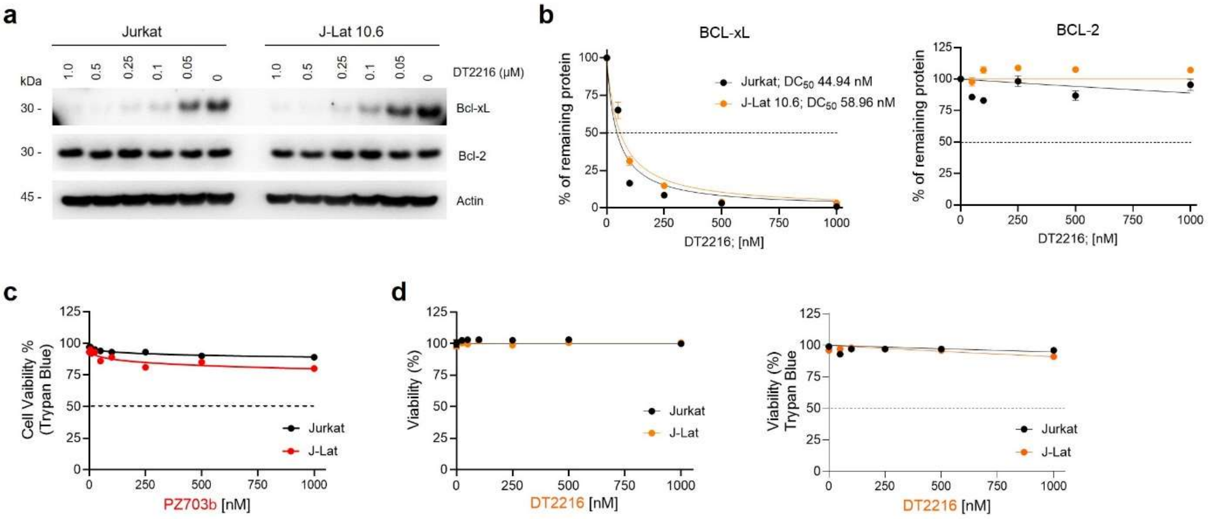


**Supplementary Fig. 2 DT-2216, a Bcl-xL PROTAC, selectively induces Bcl-xL degradation without cytotoxicity in Jurkat and J-Lat cells. a**, **b** DT-2216 efficiently degrades Bcl-xL but not Bcl-2 proteins in Jurkat and J-Lat 10.6 cell lines after 24 hours of treatment with increasing concentrations. DC50 represents the drug concentration required for 50% protein degradation. Actin was used as a loading control in immunoblot analyses. The density of untreated samples was set to 100% for protein levels; treated samples were first referenced to Actin and then normalized to untreated to calculate the reduction percentage. **c**, **d** MTS and trypan blue exclusion assays were performed in Jurkat and J-Lat cells treated with increasing concentrations of PZ703b (**c**) or DT-2216 (**d**) for 24 hours.

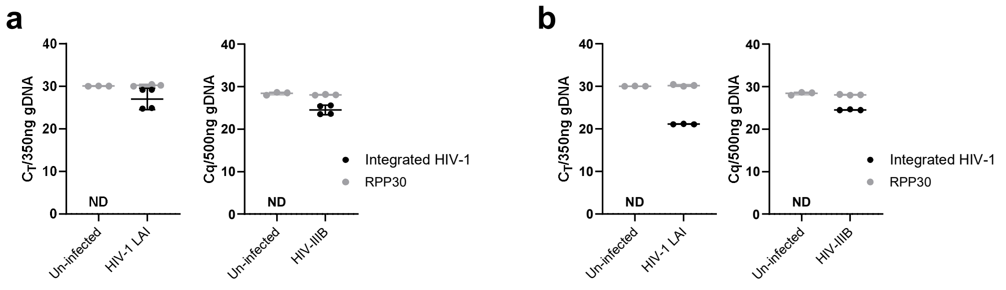


**Supplementary Fig. 3 Confirmation of HIV-1 Latency in the $T_{CM}$ model of latency with ARTs**. **a** RAL and ENF, or **b** RAL and DRV. Integrated HIV-1 DNA displayed average Ct values of 20 and 25, corresponding to approximately 300 to 3,000 copies of HIV-1 latent proviruses in the HIV-LAI and HIV-IIIB latently infected $\mathbf{T_{CM}}$ cells, respectively[54].

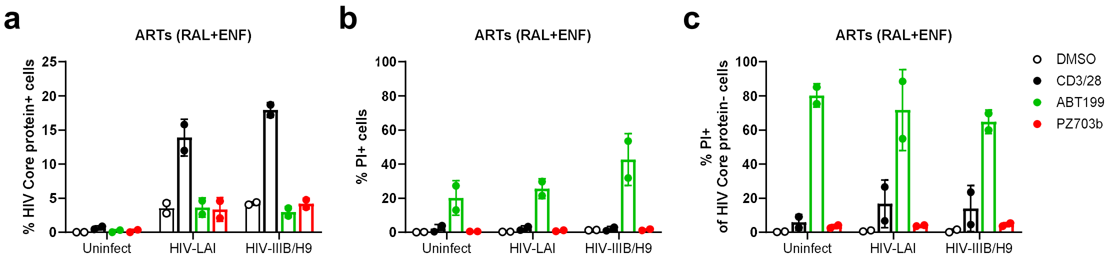


**Supplementary Fig. 4 The latency reversal and killing activities of PZ703b and ABT199 in the presence of a combination of integrase inhibitor and fusion inhibitor in $T_{CM}$ model of latency.** Naive CD4+ T cells were isolated from the peripheral blood mononuclear cells (PBMCs) of two healthy donors and established in a latent state by infection with HIV-LAI or HIV-IIIB/H9, or left as uninfected controls, as described in the Methods section.

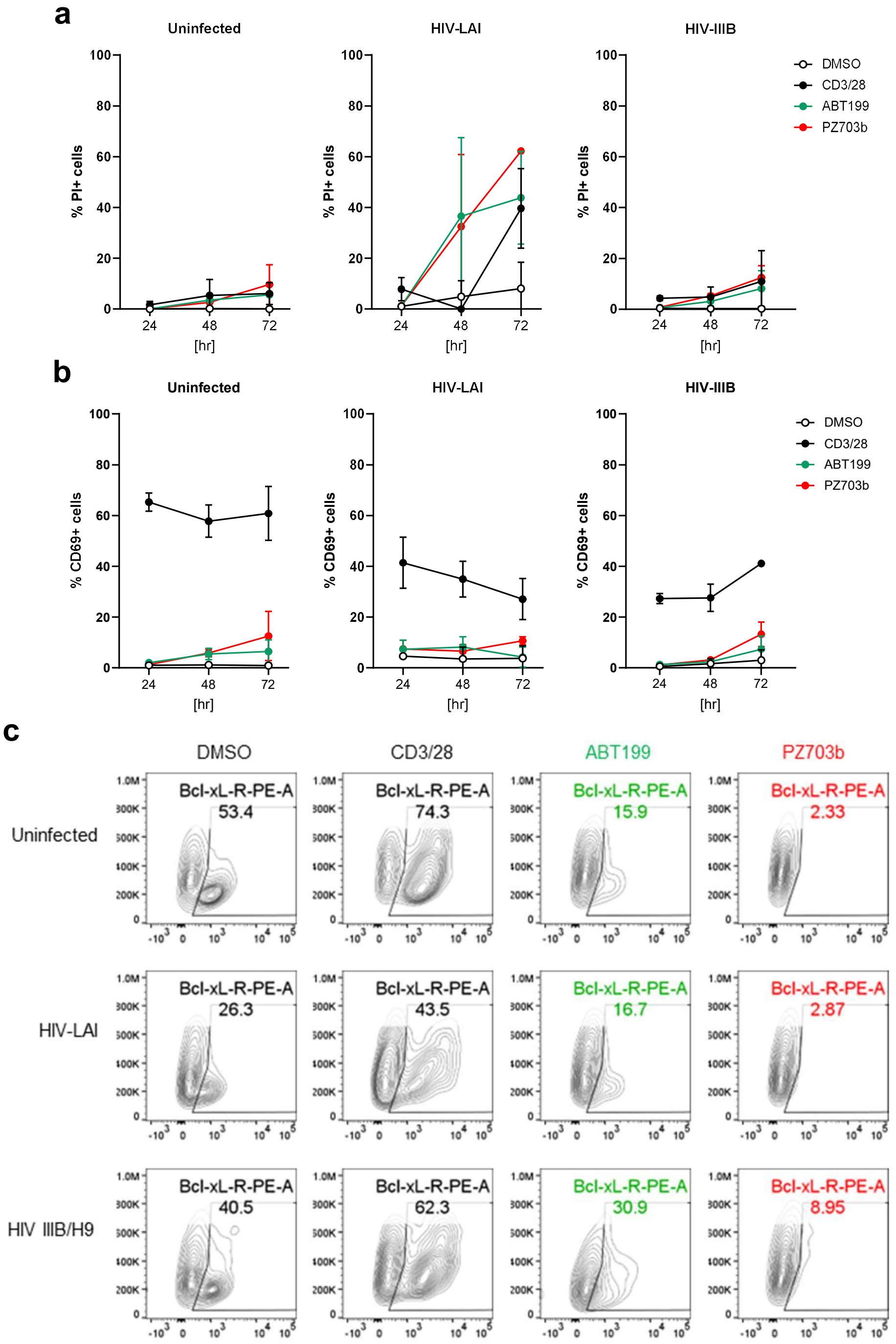


**Supplementary Fig. 5 Effects of PZ703b and ABT199 on Cell Toxicity, Global Activation, and BCL-xL Degradation in the $T_{CM}$ model of latency**. The frequency of propidium iodide-positive (PI+) cells as in **a**, CD69-positive (CD69+) cells as in **b**, and BCL-xL expression levels as in **c** following treatment of latently infected cells with DMSO, anti-CD3/CD28 antibody-conjugated beads, ABT199 (1 μM), or PZ703b (0.5 μM) for 24, 48, and 72 hours is presented as mean percentages ± SD from two biological replicates.

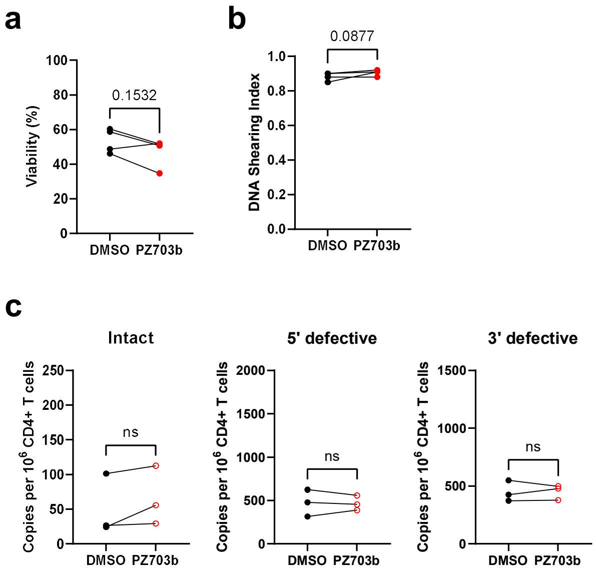


**Supplementary Fig. 6 shows a, b** the cell viability and shearing index of ex vivo CD4+ T cells and their corresponding DNA samples. **c** results of the IPDA assay of ex vivo reservoirs from HCV-negative donors treated with PZ703b

**Supplementary Table 1.** Clinical features of the four participants with HIV whose peripheral blood mononuclear cells were used in the ex vivo study.

| ID | Age | Sex | HCV | Height (cm) | Weight (kg) | HIV Regimen | CD4 T cells | Viral load |
|---|---|---|---|---|---|---|---|---|
| 1 | 49 | Female | + | 167.9 | 105.7 | Abacavir, lamivudine, darunavir/ritonavir | 953 | <50 |
| 2 | 57 | Female | + | 157.5 | 79.4 | Tenofovir disoproxil fumarate, nelfinavir | 794 | <50 |
| 3 | 65 | Female | + | 147.6 | 56.5 | Tenofovir disoproxil fumarate, lamivudine, lopinavir/ritonavir saquinavir | 1144 | <50 |
| 4 | 57 | Female | + | 157.5 | 79.4 | Tenofovir, lamivudine, nelfinavir | 794 | <50 |
